# Computing the mechanism of *α*-helix to *β*-sheet transition in proteins using the finite temperature string method

**DOI:** 10.1101/2022.07.21.500930

**Authors:** Avijeet Kulshrestha, Sudeep N Punnathanam, K Ganapathy Ayappa

## Abstract

The transition of an *α*-helix to a *β*-sheet in proteins is among the most complex conformational changes seen in bio-molecular systems. Currently, it is difficult to study such protein conformational changes in a direct molecular dynamics simulation. This limitation is typically overcome using an indirect approach wherein one computes the free energy landscape associated with the transition. Computation of free energy landscapes, however, requires a suitable set of collective variables that describe the transition. In this work we demonstrate the use of path collective variables [*J. Chem. Phys*. **126**, 054103 (2007)] and combine it with the finite temperature string (FTS) method [*J. Phys. Chem. B* **109**, 6688-6693 (2005)] to determine the molecular mechanisms involved during the structural transition of the mini G-protein from an *α*-helix to a *β*-hairpin. The transition from the *α*-helix proceeds via unfolding of the terminal residues, giving rise to a *β*-turn unfolded intermediate to eventually form the *β*-hairpin. Our proposed algorithm uses umbrella sampling simulations to simulate images along the string and the weighted histogram analysis to compute the free energy along the computed transition path. This work demonstrates that the string method in combination with path collective variables can be exploited to study complex protein conformational changes such as a complete change in the secondary structure.

## I. INTRODUCTION

Protein conformational changes involving its secondary and tertiary structures play an important role in many biological processes^1^. The secondary and tertiary structures of a protein are held together by hydrogen bonds, salt bridges, hydrophobic interactions, and disulfide bonds^2–5^. At room temperature conditions, these bonds are strongly affected by the thermal motion of the atoms of the proteins as well as those present in its surroundings including solvents and other macromolecules. Thus, the stability of various protein conformers is determined by an interplay between the interaction energy and entropy; in other words by its free-energy^6^. Simulations of conformational changes in proteins through equilibrium molecular dynamics are challenging because the timescales involved in these processes range from a few micro to milliseconds^7,8^. These timescales are much longer than those accessible in a typical molecular dynamics simulation. Instead, these conformational changes can be modeled as a barrier crossing event from one free-energy minimum to another. The dynamics then resemble a random-walk on a free-energy landscape, which can be described by a Fokker-Planck equation. The solution of the Fokker-Planck equation requires the knowledge of the underlying free-energy landscape, which makes it imperative to have techniques that can reliably compute the transition path of protein conformational changes and the free energies associated with the path.

Since it is practically impossible to follow the positions of all the atoms, the process is usually described by a few relevant variables called ‘collective variables’ (CVs). The free energy landscape is then conveniently described in terms of these CVs. There exist in literature, a variety of methods for computing free energy landscapes of various systems undergoing the physical process. As explained by Branduardi, Gervasio, and Parrinello ^9^, these methods broadly fall into two categories. In the first category, methods such as umbrella sampling, metadynamics, steered molecular dynamics and adaptive force bias compute the entire free energy landscape described by the CVs^10–14^. The second category is path-based methods such as nudged elastic band^15^, string method^16^, etc., which can compute the transition path in a highly efficient manner if the initial and final states are known. These are also known as the chain of state methods and use a set of interpolating images to represent the transition path. Depending on the system and the algorithm, the computed transition path could be the minimum energy path (MEP)^16^, minimum free-energy path (MFEP)^17^, the principal curve^18^, or the most probable transition pathway (MPTP)^19^. One of the earliest such methods is the plain elastic band method^20^. In this method, each image is connected to its two neighboring images through springs. The main drawback of this method is the so-called corner-cutting that occurs when a high value of the spring constant is used. A high value of the spring constant causes the images to be pulled away from the MEP/MFEP by the component of the spring force perpendicular to the path. The nudged elastic band (NEB) method^15^ was developed to circumvent this problem. In the NEB approach, the parallel component of the spring force is used to keep the images on the path from falling into the free energy minima and only the perpendicular component of the true force is used to evolve the path towards the MEP or the MFEP. There exist many variations of the original NEB method such as the adaptive NEB^21^, the doubly NEB^22^, etc. A different method derived from the plain elastic band approach is the string method, which uses a simple reparameterization of the images along the path to avoid movement of the images towards the minima and simultaneously retain images at equal distances on the path^16^. The original implementation^16^ is called the zero temperature string (ZTS) method and it computes the MEP on an energy landscape. For finite temperatures, two variants of the string method were developed, namely, the string method in collective variables (SMCV)^17^ that computed the MFEP and the finite temperature string method (FTS)^18,23^ that computed the principal curve. The MFEP curve is parallel to the mean force on the free energy landscape in an appropriate metric. Since it depends only on the local characteristics of this landscape, the MFEP may still miss important non-local features of the free energy landscape in the directions perpendicular to itself^23^. In contrast, a principal curve is such that its position coincides with the average position within the hyperplanes perpendicular to the curve. Hence it is a more natural candidate to explain the mechanism of the transition since its location depends more globally on the features of the underlying free energy landscapes^23^.

One of the challenges in studying conformational changes in molecules is the identification of a suitable set of CVs that adequately represents the conformation change. In general, this is not a trivial task and the choice of CVs depends strongly on the system/molecule being studied. This is especially true for protein conformational changes which involve many complex changes in the secondary structure of the protein^8^. To this end, the path collective variables (PCVs) developed by Branduardi, Gervasio, and Parrinello ^9^ form a convenient set of orthogonal collective variables that can potentially be applied to characterize the conformational change for any molecule. Given that one is usually interested in two end states of the system, PCVs offer a convenient framework to define a sequence of reference structures that connect the two end states in the transformation. PCVs have been applied to study transitions and free energy barriers for a wide variety of systems. They have been used to build a neural network based reaction potential for urea decomposition^24^, study enzymatic reactions^25,26^, obtain the free energy landscape for methane and isocyanic acid conversion into methyl isocyanate^27^ as well as a few pharmacologically relevant problems^28,29^. Using metadynamics simulations in an all-atom framework, PCVs have been successfully applied to study the free energy landscape associated with large conformational changes in kinases^30,31^. Recently Cuendet *et al*. ^32^ used the PCVs to compute the free energy difference between *α*-helix and *β*-hairpin conformations of deca-alanine in solution, however, the free energy landscape was not evaluated.

There are a few studies where the string method has been used to evaluate conformational free energies of proteins^8,23,33–36^. In the work by Ovchinnikov and Karplus ^8^ the free energy of the transition in a G-protein between the *α*-helix to *β*-hairpin states has been studied in implicit solvent and Song and Zhu^36^ combined the string method with umbrella sampling to study side chain flipping in a bacterial transporter using replica exchange molecular dynamics. Vanden-Eijnden and Venturoli ^23^ used a highly coarse grained model of a protein with the string method for studying the conformational transition of the nitrogen regulatory protein C and Matsunaga *et al*. ^35^ used the string method in combination with the multistate Bennett acceptance ratio to study the adenylate kinase enzyme transition in explicit solvent.

In this work, we compute the transition pathway involved during the structural change from an *α*-helix to a *β*-hairpin in a mini-G protein. The FTS method is used to compute the transition pathway with the free energy landscape determined through umbrella sampling simulations. This is a highly complex conformational change that involves a complete change in the secondary structure of the protein. Such a change has only been previously studied by Ovchinnikov and Karplus ^8^. In that study, they had to use 107 collective variables to characterize the free energy landscape. In this work, we demonstrate that it is possible to characterize the free energy landscape involved in this transition using PCVs; thus reducing the number of collective variables to just two in number. In spite of such a substantial reduction in the number of collective variables, the molecular mechanism of this transition along the computed pathway is qualitatively similar to that discovered by Ovchinnikov and Karplus ^8^. All the molecular dynamics simulations needed for this computation can be performed using GROMACS-2018.6^37^. The use of the PCVs in the simulations can be done through the PLUMED package^38^.

## II. METHOD

### A. Path collective variables (PCVs)

The PCVs were first introduced by Branduardi, Gervasio, and Parrinello ^9^ and is defined for any *N* particle system with positions (**r**_1_, **r**_2_, …, **r**_*N*_) ≡ **R** undergoing a conformational change. We first consider a sequence of *n* reference structures, {**R**^(1)^, **R**^(2)^, …, **R**^(*n*)^} of the molecule under consideration, connecting two endpoints of the conformational change. The two PCVs, *S* and *Z*, are then defined as

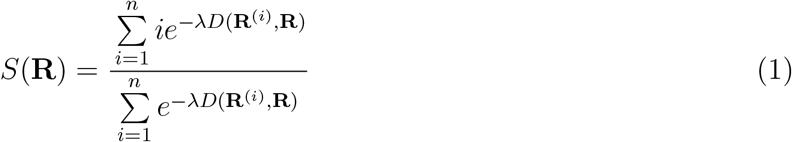

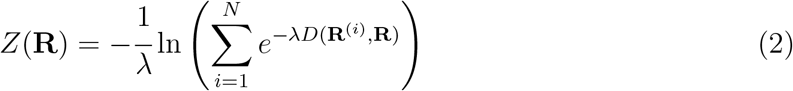

where *S* is a measure of the distance traveled from one endpoint to the other and *Z* denotes the movement in the perpendicular direction. The idea of the *S* and *Z* for a linear reference path in the Cartesian coordinate system is plotted in supplementary Figure S1. *D*(**R**^(*i*)^, **R**) is the distance between the dynamic instantaneous coordinates of the molecule **R** and the reference structures coordinates **R**^(*i*)^. Although there can be several ways to define the distance, in this work we define it to be the mean square displacement (MSD), i.e.,

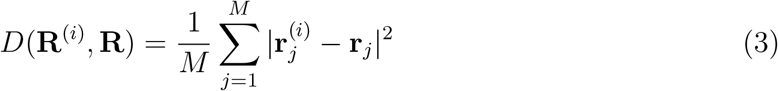

where *M* is the number of sites in the molecule specified *a priori*. As *λ* → ∞, *S* takes integer values from *i* = 1 (if instantaneous coordinates are at the first reference structure) to *i* = *n* (if instantaneous coordinates of the system are at *n*^*th*^ reference structure). For other values of *λ*, the PCVs phase-space merely changes the location of instantaneous structures. In practice, the value of *λ* is chosen to be approximately inverse of the MSD between two consecutive reference structures.

### B. Finite Temperature String Method (FTS)

The transition path computed using the FTS method is the principal curve. In the original formulation^18^, the principal curve was computed iteratively by performing simulations that were restricted to the perpendicular hyperplanes. Later, the same authors presented a simplified algorithm where the hyperplanes were replaced by Voronoi cells, whose respective generating points are the discretization points or images along the string^23^. In each iteration, the molecular dynamics simulation for each image is restricted to its respective Voronoi cell by implementing a collision rule at the boundaries of the cell. The generating points are then moved to the average position within the cells till convergence of the string is achieved. The advantages of the modified algorithm are (i) simplified calculation of the free energy along the transition pathway and (ii) easy extension to computing transition pathway in collective variable space.

In this work, instead of restricting an image to its Voronoi cell, we use umbrella sampling simulations for each image. The following harmonic function is used as the biasing potential, Ψ, for the umbrella sampling simulations.

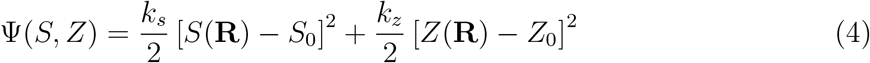

During each simulation, we compute the average location of the image in the collective variable space, i.e. ⟨*S*⟩ and ⟨*Z*⟩. The locations of the images are then updated to these average locations and then reparameterized as per the original string method^18^. The above iterative process is continued till convergence of the string is attained. A schematic representation of this algorithm is shown in Fig. 1. A significant advantage of this algorithm is the avoidance of the estimation of the complex M-tensor which accounts for the curvilinear nature of the collective variables^23^. In some studies^39^, this tensor is assumed to be unity for sake of computational efficiency. In this method, we implicitly assume that the biasing potential does not preclude the sampling of all the relevant local features for the determination of the principal curve. The free energy along the converged string was then computed from these umbrella sampling simulations using the WHAM algorithm^40^.

**FIG. 1.**
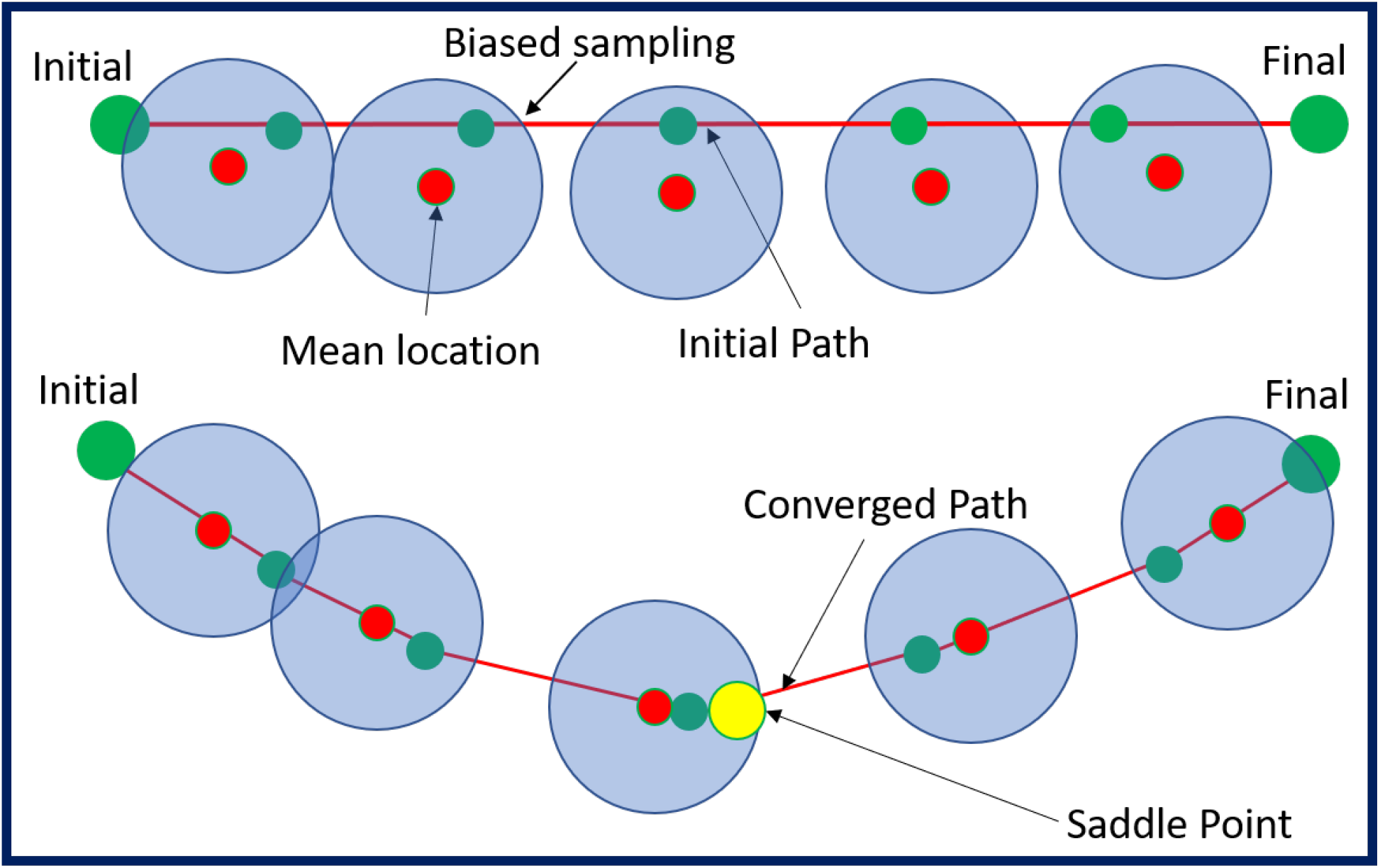
Schematic representation of the evolution and convergence of the string using the FTS method. The red line represents the string. The green circles represent the current location of the images along the string. The large blue circle represents the phase space sampled for each image during the umbrella sampling simulation and the red circle represents the average position in each simulation. The yellow circle represents the saddle point in the free energy landscape.

## III. METHOD VALIDATION

We validated our method by studying the conformational change of the alanine-dipeptide in a vacuum and comparing the computed transition path with the one reported in Ren *et al*. ^41^. The conformational change of alanine-dipeptide has been well studied and serves as an excellent test for any method that computes transition pathways. Alanine-dipeptide can exist in two main conformers, namely *C*_7*eq*_ and *C*_*ax*_, which differ from each other in the values of the backbone dihedral angles *ϕ* and *ψ* (see Fig. 2A). The state *C*_7*eq*_ is further split into two sub-states (*C*_7*eq*_ and 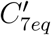, which are separated by a small barrier. In this work, the alanine-dipeptide was modeled using the CHARMM22 forcefield^42^. The temperature of the system was maintained at 272 K using a velocity rescale thermostat^43^. The string representing the transition from *C*_*ax*_ to 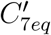 consists of 27 images. Umbrella sampling simulations with the harmonic bias potential were performed for each image and the location of the image was progressively updated according to mean values of *ϕ* and *ψ*. At the start of every iteration, the positions of the images in the string were reparameterized such that they are equidistant from each other. The process was continued till the positions of the images converged (see Fig. 2B). The converged transition path is shown in Fig. 2C. The free energy change along the string was computed from the results of the umbrella sampling simulations using the WHAM algorithm^40^(Fig. 2C). We have determined the free energy barrier to be 16 *k*_*B*_*T* for transition from *C*_*ax*_ to *C*_7*eq*_ (see Fig. 2D). Since this is small single molecule system, we further validated our results by computing the free energy landscape using well-tempered metadynamics simulations^44^. Metadynamics simulation was performed on *ϕ* and *ψ* collective variables where the initial Gaussian height was 1.2 kJ/mol, Gaussian width was 0.35 radian for both CVs, and Gaussians were deposited every 1 ps, and the bias factor was 6.0. A convergence check of the metadynamics is given in supplementary Figures S2A and S2B. The computed free energy landscape is shown in Fig. 3. One can clearly observe that the computed transition path passes through the saddle point in the free energy landscape. We also compare our principal curve with the transition path computed by Ren *et al*. ^41^ in Fig. 3 where the literature values are scanned from the previously published figure. The two results show excellent agreement with each other, thereby validating our method for computing the principal curve. Committor distribution test was done by considering 200 random states at the saddle point. Subsequently, 5000 trajectories of 6 ps were launched from each random states, and committor distribution *P*_*Cax*_ for trajectories that fall into *C*_*ax*_ stable state was calculated. Committor distribution shows a peak at 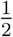 (Figure S2C) that reinforces the reliability of the obtained transition state ensemble.

**FIG. 2.**
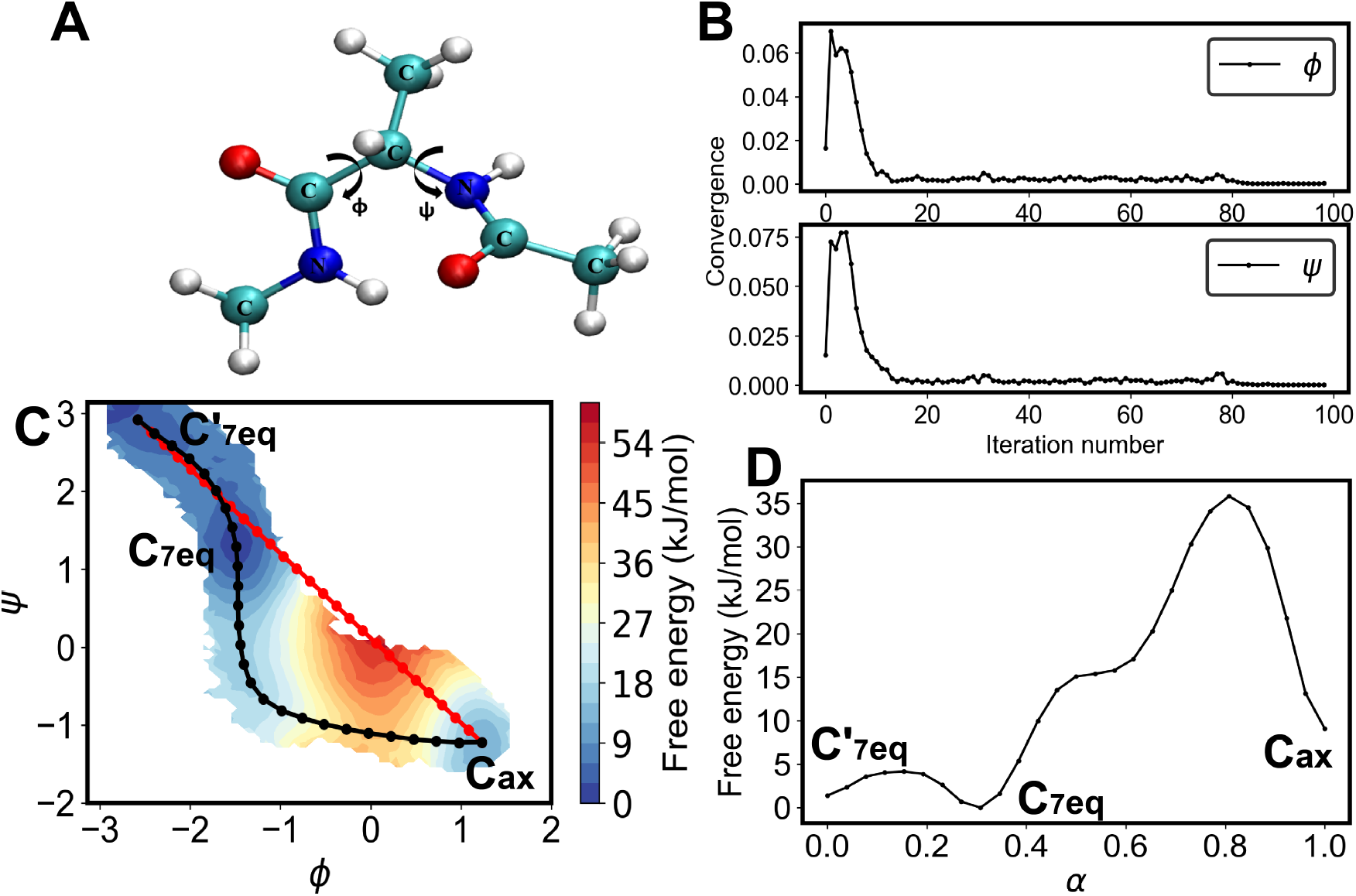
Alanine-dipeptide. (A) Depiction of the two dihedral angles in alanine-dipeptide. (B) Convergence of *ϕ* and *ψ* during the string evolution. (C) Free energy landscape of alanine-dipeptide in vacuum obtained from the WHAM analysis. A linear initial guess (red) of the path and the converged principal curve (black) are depicted. (D) Free energy profile as a function of the parameterized distance ‘*α*’ between the ends of the string.

**FIG. 3.**
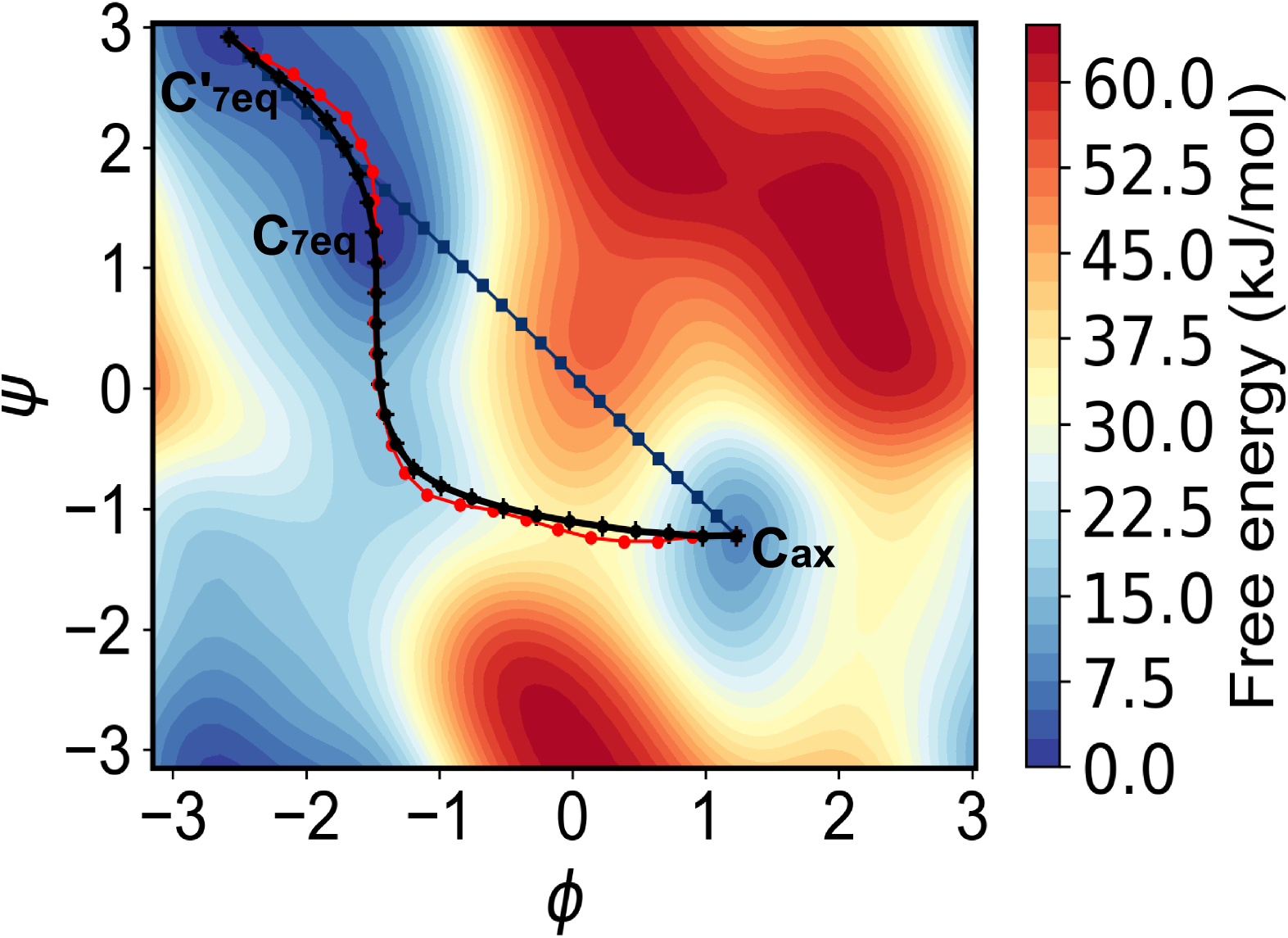
Alanine-dipeptide. Free energy landscape produced by well tempered metadynamics along with the initial string (blue color), converged string (black color), and the transition path reported in the literature (red color)^41^.

## IV. *α*-HELIX TO *β*-HAIRPIN TRANSITION IN MINI G-PROTEIN

G-protein is an immunology binding protein expressed in group C and G Gram-positive streptococcal bacteria. It is a Y-shaped protein (antibody, Ab) used by the immune system to neutralize pathogens such as pathogenic bacteria and viruses^45^. The G-protein structure consists of 56 residues and has a molecular weight of 6.20178 kDa. The last 16 residues (41-56) with the sequence of *GEWTYDDATKTFTVTE* of this protein form a *β*-hairpin (Fig 4B). Mini G-protein *β*-hairpin is a stable conformer^46^ and it has much of the complexity of realistic proteins, and is therefore used in literature to study the folding free energy landscape^47,48^ and the conformational change. Ovchinnikov and Karplus ^8^ studied the conformational change of this mini G-protein from its stable *β*-hairpin X-ray structure to an *α*-helix using the FTS method where the *α*-helical conformation was generated from default internal coordinate entries of CHARMM with backbone dihedrals set to −57^*o*^ and −47^*o*^. The collective variables needed to define the string images were the coordinates of all the heavy atoms except for certain heavy side-chain atoms. The protein interactions were modeled using the CHARMM36^49^ forcefield and the effects of surrounding water molecules were accounted using an implicit FACTS solvation model^50^. This was one of the earliest demonstration of the utility of the FTS method for computing the transition paths and associated free energy profiles for a conformational change involving a complete change in the secondary structure of the protein. They found many reaction barriers in the free energy profiles for each of the computed transition paths.

**FIG. 4.**
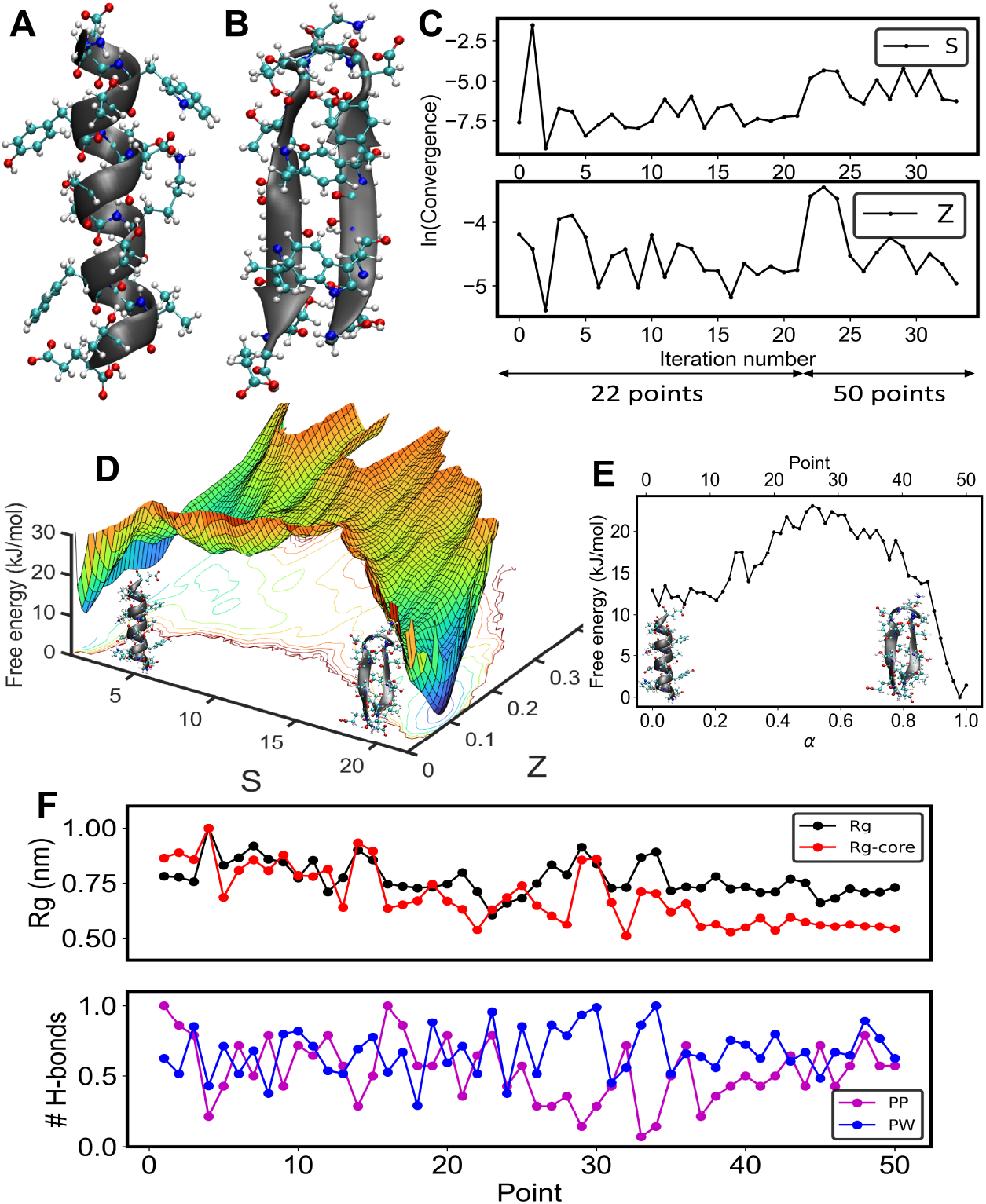
G-protein. (A) G-protein *α*-helix state. (B) G-protein *β*-hairpin state. (C) Convergence of collective variables *S* and *Z* during the evolution of the string. (D) 3D surface plot of the free energy landscape indicates a rugged landscape with multiple metastable states of the order of *kT* illustrating the transition from the *alpha*-helix (low *S*) to the *beta*-hairpin state (high *S*). (E) Free energy profile versus the parameterized distance ‘*α*’ from *α*-helix state to *β*-hairpin state on the string. (F) Changes in the radius of gyration of protein ‘*R*_*g*_’, and the side chain of the core hydrophobic residues (W43, Y45, F52, and V54) ‘*R*_*g*_-core’ along the string. The number of intra-protein (PP) hydrogen bonds and hydrogen bonds between protein and the water molecules (PW). Both the *R*_*g*_ and hydrogen bond values on the path are normalized with their respective maximum values, *R*_*g*_ 1.07 nm, *R*_*g*_-core 1.00 nm, H-bonds (PP) 14, H-bonds (PW) 93, on the path.

In this work, we revisit this study of the *α*-helix to *β*-hairpin transition of the mini-G protein with two significant differences: (i) the number of collective variables was substantially reduced to just two through the use of PCVs, and (ii) the effects of the surrounding solvent molecules were modeled explicitly through the TIP3P model for water. The protein interactions were modeled with the CHARMM36^49^ forcefield. Similar to the study in Ref. 8, the simulations were performed at 300 K. The *β*-hairpin structure of the mini G-protein was taken from the protein data bank (PDB 2GB1). This structure was first energy-minimized in a vacuum and then solvated with TIP3P water. Following this, the system was equilibrated in two steps. In the first step, an NVT simulation was performed at 1 ns with 3094 molecules of water inside a cubic simulation box having a box length of 4.6 nm. In the second step, an NPT simulation was performed for 1 ns at a pressure of 1 atm. In both of these steps, a position restraint was applied to the heavy atoms of the protein to maintain the *β*-hairpin structure. The isotropic pressure control was achieved using the Parrinello-Rahman method^51^ with a time constant of 2 ps, and the temperature was controlled using the velocity rescale thermostat^43^ with a time constant of 0.1 ps. The long-range electrostatic interactions were treated using the particle mesh Ewald (PME) method^52^ with a real space cut-off of 1.0 nm. Three-dimensional periodic boundary conditions were applied to eliminate boundary effects. All bonds were constrained using the LINCS constraint, which allowed a larger time step of 2 fs^53^. The solvent and the protein were coupled to a temperature bath. Pressure was kept constant at 1 bar using the isothermal compressibilities of *κ*= 4.5 × 10^*−*5^ bar^*−*1^. The equilibration of the system was monitored by evaluating the root mean squared deviations (RMSD) for the protein. In 100 ns of simulation, *β*-hairpin shows an average root mean square deviation (RMSD) of 2 Å. The *α*-helix structure of the mini G-protein was created with the help of Avogadro^54^ with the backbone dihedrals set to values −57^*o*^ and −47^*o*^ (Fig 4A) similar to Ref. 8. Later this structure was minimized in a vacuum and then solvated with water. The solvated structure was then equilibrated using the same protocol as that used for the *β*-hairpin structure. However, in a restraint free simulation, the end-terminus of the *α*-helix unfolds within 40 ns and the protein loses its helicity in 100 ns. Given this situation, we used the initial helix structure as an endpoint of the string (Fig 4A). Unrestrained simulation results are shown in supplementary Fig. S3.

The reference path for the PCVs consisted of a set of 22 structures generated by linear interpolation between the *α*-helix and the *β*-hairpin structures. For the interpolation, three different cases of the atom coordinates were considered: (i) only backbone atoms coordinates, (ii) all heavy atom coordinates, and (iii) coordinates of all atoms including hydrogen. To make an appropriate choice between these three sets, we performed pulling MD simulations from the *α*-helix structure to the *β*-hairpin. The external force for these pulling simulations was defined with respect to the PCVs. After the protein structure attained the PCVs values close to the *β*-hairpin structure, all restraints were removed and an unbiased MD simulation (200 ns) was performed to check its stability. Unbiased simulation results are given in supplementary Fig. S4 where only the heavy atom and all-atom coordinates form a stable *β*-hairpin. However, the analysis of the secondary structural change with time indicates that the *β*-hairpin structure was most stable when all-atom coordinates were used. The main reason for this result was the incomplete formation of hydrogen bonds when only backbone and heavy atoms are used. Since the formation of hydrogen bonds is necessary to impart stability to the *β*-hairpin structure, we conclude that including the coordinates of the hydrogen atoms refines the definition of the reference path. The RMSD between two consecutive structures in the reference path is 0.048 nm, which is well below 1 Å as recommended by Branduardi, Gervasio, and Parrinello ^9^ Accordingly, in the definition of PCVs, we use a value of *λ* = 998.26 nm^*−*2^ which is equal to 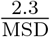

The initial path for the string method was generated by pulling the *α*-helix towards the *β*-hairpin. During these simulations, the value of *Z* was restrained to be close to zero and the value of *S* was changed progressively from 1 to 22. The value of the force constant during these pulling simulations varied between 1000 and 10000 kJ/mol. The transition path was represented by a set of 22 images and the principal curve was computed iteratively by performing umbrella sampling simulations for 2 ns at each image with the values of both *k*_*s*_ and *k*_*z*_ set to 10 kJ/mol. After 23 iterations, we did not observe any further evolution of the transition path. The number of points on the string was further increased to 50 to capture the transition in detail and improve the sampling along the path. Subsequently, 13 more iterations were performed to evolve the string. The location of converged strings defined by 22 and 50 points remains within a few *k*_*B*_*T*; however, the string with 50 points indeed captured the ruggedness with greater detail (see Fig. S5). To further improve the precision of the mean estimation and improved convergence of the free energy profile, an additional iteration was performed with 50 ns simulations at each image. The sum of standard error of mean reduced from 4.4 × 10^*−*4^ to 1.2 × 10^*−*4^ for the *S* variable and 2.2 × 10^*−*5^ to 9.4 × 10^*−*6^ for the *Z* variable.

After the string convergence, the free energy landscape under the principal curve was calculated using the WHAM^40^ technique from data obtained in the umbrella sampling simulations. The convergence of the free energy calculation along the path is analyzed by the changes in the free energy profile along the string with the increase in sampling as shown in Fig. S5A. Figure 4D shows the free energy landscape and one can clearly observe that the landscape is very rugged similar to the observations by Ovchinnikov and Karplus ^8^. Another study by Zhou, Berne, and Germain ^48^ had also shown that the free energy landscape of folding of *β*-hairpin is rugged at low temperatures, becoming smoother only at higher temperatures. The free energy profile as a function of the parameterized distance ‘*α*’ (in the collective variable space) between the two ends of the string is shown in Fig 4E. The maximum barrier is 23.03 kJ/mol which is lower than that of the barrier computed by Ovchinnikov and Karplus ^8^ where they found barrier heights in the range of 40-70 kJ/mol for the different transition paths. Our calculations differ from that of Ref. 8 through the use of an explicit model for the surrounding water molecules and the lower free energy barrier obtained from our simulations highlights the importance of the role played by explicit water during changes in the secondary structure of the protein.

An analysis of the structural changes seen during the transition emphasizes the role of hydrogen bond formation and the radius of gyration, *R*_*g*_ (Fig. 4F), where the structures on the path were generated by carrying out a short MD simulation of 1 ns with a high spring constant on PCVs along the converged path. For each biased simulation on the path, the protein coordinates were taken from the final configuration from the string evolution. Since *R*_*g*_ (scaled with the maximum value on the path) shows significant variation across the transition, the mechanism for the conformational change appears to be driven by hydrophobic effects. The *R*_*g*_ of the side chain of *β*-hairpin core hydrophobic residues (W43, Y45, F52, and V54), *R*_*g*_-core shows higher values in the helical state and approaches to a lower value as the peptide evolves towards the hairpin state. In general, we observe that an increase in *R*_*g*_ is concomitant with a decrease in the intra-protein (PP) hydrogen bonding, reflecting changes in secondary structure or increased disorder during the intermediate states of the transition. However, above point 37 (Fig 4F) on the string, *R*_*g*_ is relatively invariant, and the *β*-hairpin stabilizes with an increasing trend observed in hydrogen bonding. This trend is observed in the structural snapshots illustrated in Fig 5. An increase in the protein-water hydrogen bonds (PW) was also observed during the unfolding of the protein, indicating the role of water molecules in the conformational change. The folding of the *β*-hairpin from an unfolded state in explicit solvent was also found to occur with a simultaneous decrease in *R*_*g*_ and hydrogen bond formation^48^.

**FIG. 5.**
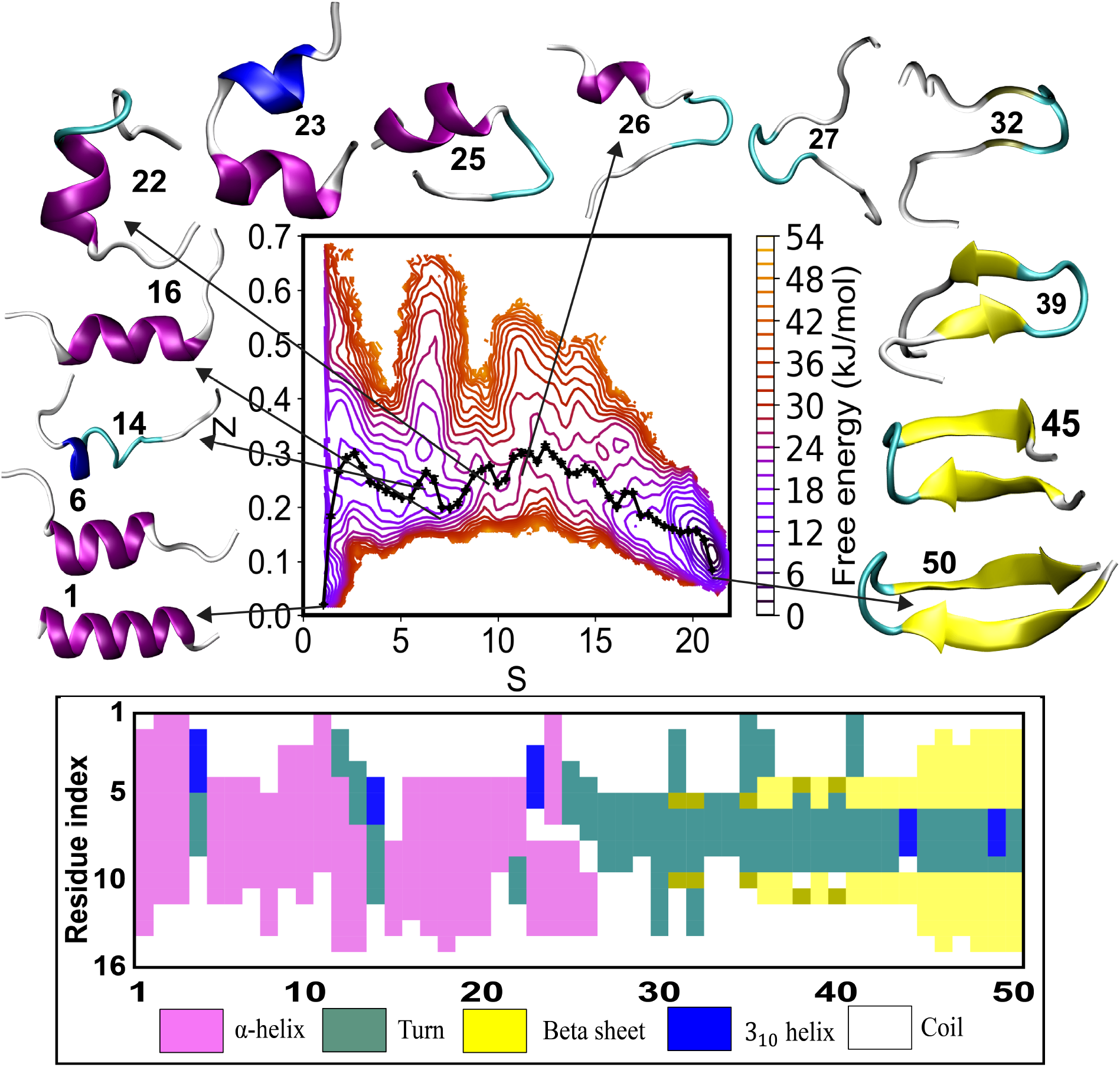
*α*-helix to *β*-hairpin transition of G-protein. Free energy surface in the *SZ*-plane along with the converged path of the string is illustrated. Snapshots that capture the key changes in secondary structure along the string are also shown. Residue-wise secondary structure changes along the 50 points on the string from a STRIDE analysis is illustrated below.

Figure 5 illustrates the changes in the secondary structure during the conformational change from *α*-helix to *β*-hairpin structure. We observe that the *α*-helix first unfolds from the C-terminus followed by progressive unfolding from both ends. However, this progression is not continuous and we observe a partially unfolded state (point 14) due to the presence of a neighboring minimum state which is partially sampled at higher Z values in the local neighborhood of the string. Further, a reduction in the free energy landscape occurs at point 16, associated with an increase in helicity. A common feature in this region of the free energy landscape is the loss of helicity at the ends of the protein and a *β*-turn formation observed at point 23. Secondary structure analysis reveals a significant loss of helical content beyond this point, following which turn-coil conformations dominate the landscape. The region where the greatest disorder is observed occurs in the vicinity of the saddle point (point 26) corresponding to the maxima in the free energy barrier (Fig. 4E) where the structure preserves the previously formed *β*-turn (points 25-27). This disordered state is followed by the formation of the *β*-hairpin with both strands coming in close proximity to eventually form the *β*-hairpin structures (points 39 - 50). This region along the string is also where the number of hydrogen bonds is found to increase, consistent with the stabilization of the *β*-hairpin (Fig. 4F). Our study supports the view that the formation of the *β*-turn precedes the simultaneous formation of the hydrogen-bonded *β*-sheet as observed by Zhou, Berne, and Germain ^48^ in an explicit solvent MD simulation. A similar mechanism had been suggested earlier in an experimental study of the *β*-hairpin formation^47^. In the implicit solvent study by Ovchinnikov and Karplus ^8^, the authors computed two separate transition paths starting from two distinct initial states. The first initial path was generated by a linear interpolation combined with zero temperature string (ZTS) method and the second one by restrained targeted molecular dynamics (RTMD). The free energy barrier from the former method was relatively higher. Additionally, the transition path computed from the RTMD trajectory had a lower mean first passage time (MFPT) and thus a higher reactive flux. The structural changes in the protein sequence computed from our work are qualitatively similar to the transition path computed from the RTMD 8. Similar to Ref. 8, our simulations reveal that the *β*-hairpin is the favored structure, the helical region is found to sample a broad space in the S-Z plane (Fig. 5) where partially unfolded helical states are observed^55^.

## V. CONCLUSIONS

This work brings together a host of well-developed methods to compute complex transition paths in large bio-molecules and proposes a new variant. These include (i) the finite temperature string method to compute the principal curve, (ii) umbrella sampling simulations for each image of the string, (ii) path collective variables for characterizing the structural changes, and (iv) the weighted histogram analysis method for computing the free energy changes along the transition path. The above protocol also avoids calculation of the M-tensor in the FTS method. We illustrate the approach by evaluating the transition path for a change in the secondary structure of a 16-residue protein sequence from a G-protein, i.e. transition from an *α*-helix to a *β*-hairpin. We observe that this transition is initiated by unfolding of the end terminal residues, followed by the formation of *β*-turn, which eventually transforms to the *β*-hairpin structure. The transition occurs through an unfolded intermediate having a free energy barrier of ∼ 12 kJ/mol from the initial *α*-helical state. This unfolded intermediate state is characterized by an increased number of protein-water hydrogen bonds with a concomitant decrease in intra-protein hydrogen bonds. Qualitatively similar mechanisms for the same transition have been proposed by Ovchinnikov and Karplus ^8^ where an implicit solvent model is used. However, the free energy barriers associated with the transitions are lowered in our work, indicating the importance of explicit solvent interactions in stabilizing intermediate states during the transitions. Our method illustrates for the first time that a reduction in the number of collective variables by using PCVs combined with string based methods can be effectively used to study free energy barriers associated with secondary structure transitions typically observed in proteins. The method is generic and can potentially be used for computing the transition mechanisms for complex biomolecules involving large time scale conformational changes of solvated proteins or even highly complex membrane triggered conformational changes in membrane proteins. The algorithm is implemented in widely used biomolecular simulations software, i.e, GROMACS, and Plumed, making it readily adaptable to a variety of biomolecular systems.

## DATA AVAILABILITY STATEMENT

The data that support the findings of this study are available from the corresponding author upon reasonable request.

## ACKNOWLEDGMENTS

The funding for this work has been provided by a grant from the National Supercomputing Mission, Government of India (Grant No. DST/NSM/R&D HPC Applications/2021/03.20). The computations were carried out using computers purchased under the Nano Mission Programme of Department of Science and Technology, Government of India (Grant No. DST/NM/NS-14/2011(C)), the Cray Supercomputer (Sahasrat) of Indian Institute of Science, Bengaluru.

## Appendix A: FTS Algorithm

Let ***θ***(*x*) = {*θ*_1_(*x*), *θ*_2_(*x*), …, *θ*_*N*_ (*x*)} be the set of *N* collective variables that is used to characterize the transition. The transition path connects two end states (typically two minima in the free energy landscape associated with the above set of collective variables). The transition path computed by the FTS method is the principal curve, which is the average location of ***θ*** in the hyperplane perpendicular to the transition path. The steps involved in the computation of the principal curve are as follows.

STEP 1: We guess an initial path containing ‘P’ images of the system in the collective variables space. An umbrella sampling simulation is performed at each point on the curve with a harmonic biasing potential centered at these points. From each of these simulations, the mean location of the collective variables is computed as follows

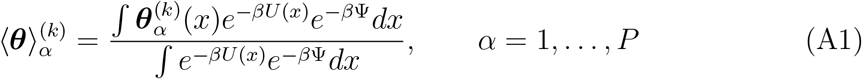

where Ψ is the harmonic biasing potential, *α* is the index of the image and *k* is the iteration number.

STEP 2: The computed mean locations from each of the above simulations form the new string at iteration *k*. Keeping the end states fixed, the locations of images constituting the string are then smoothened out using the following equation.

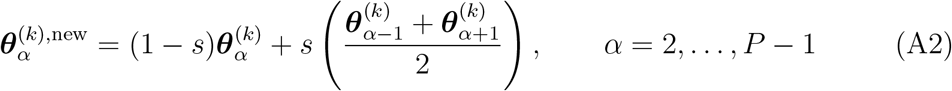

where *s* is a smoothening parameter and set equal to 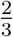 in our calculations. This essentially averages out the intermediate points on the string for every three consecutive points.

STEP 3: The locations of the new images on the string are then redistributed by enforcing equal arc length parameterization. The arc lengths are computed via linear interpolation.

STEP 4: The convergence of the string is determined by computing the quantity, *C*, as follows.

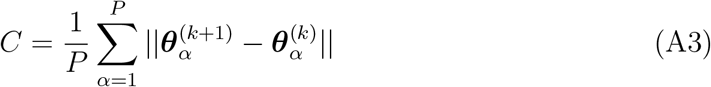

Steps 1 to 3 are repeated till the value of *C* is below the specified tolerance level.

