## supporting information for "Computing the mechanism of *α*-helix to *β*-sheet transition in proteins using the finite temperature string method"

### Supplementary information for the manuscript titled “Computing the mechanism of $\alpha$-helix to $\beta$-sheet transition in proteins using the finite temperature string method”

Avijeet Kulshrestha, Sudeep N Punnathanam,* and K Ganapathy Ayappa*

*Department of Chemical engineering, Indian Institute of Science, Bangalore, India - 560012*

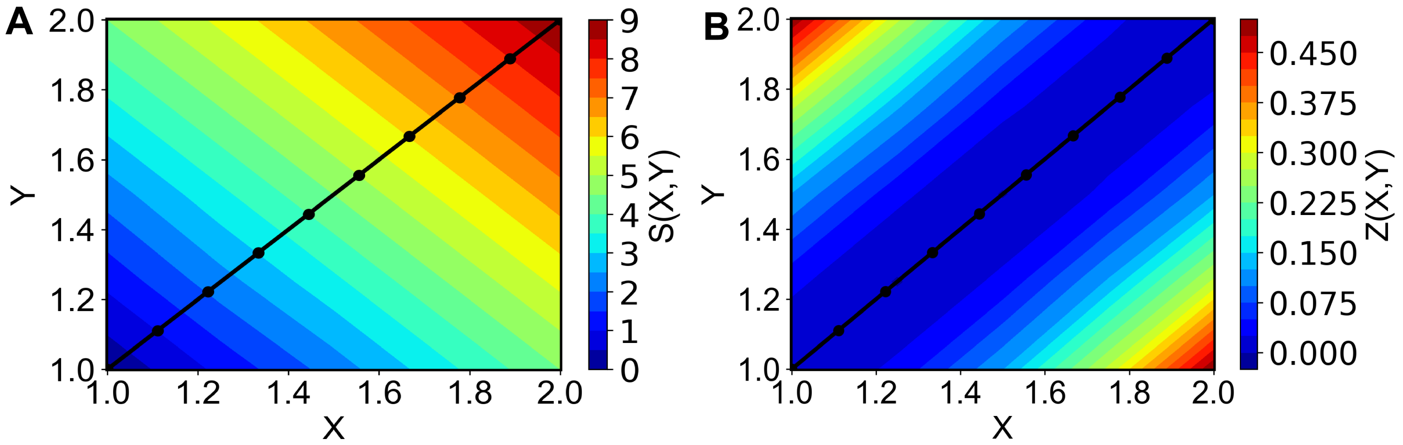


Figure 1: **Path collective variables** Projection of path collective variable (A) S and (B) Z in the 2D Cartesian coordinates system where the reference path is a linear set of ten coordinates.

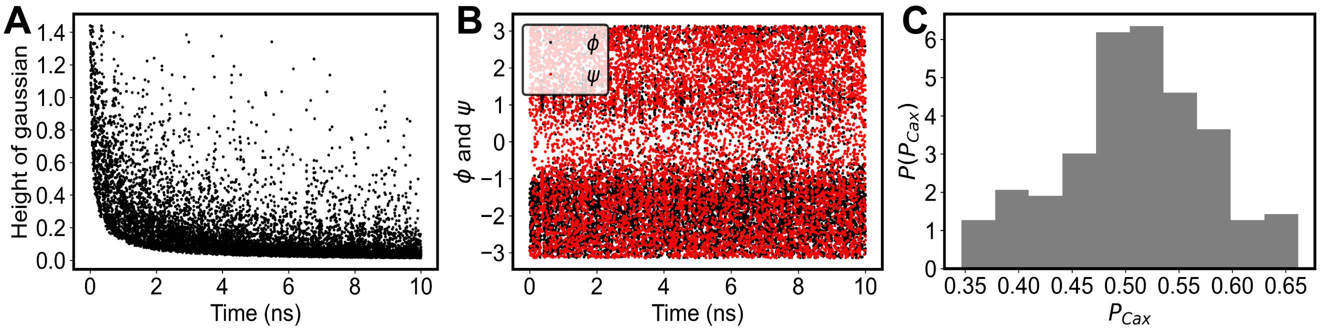


Figure 2: **Alanine-dipeptide** (A) Change in the Gaussian height during metadynamics simulations, (B) $\phi$ and $\psi$ sampling during metadynamics, and (C) Committor distribution with a committor value peaked at $\frac{1}{2}$.

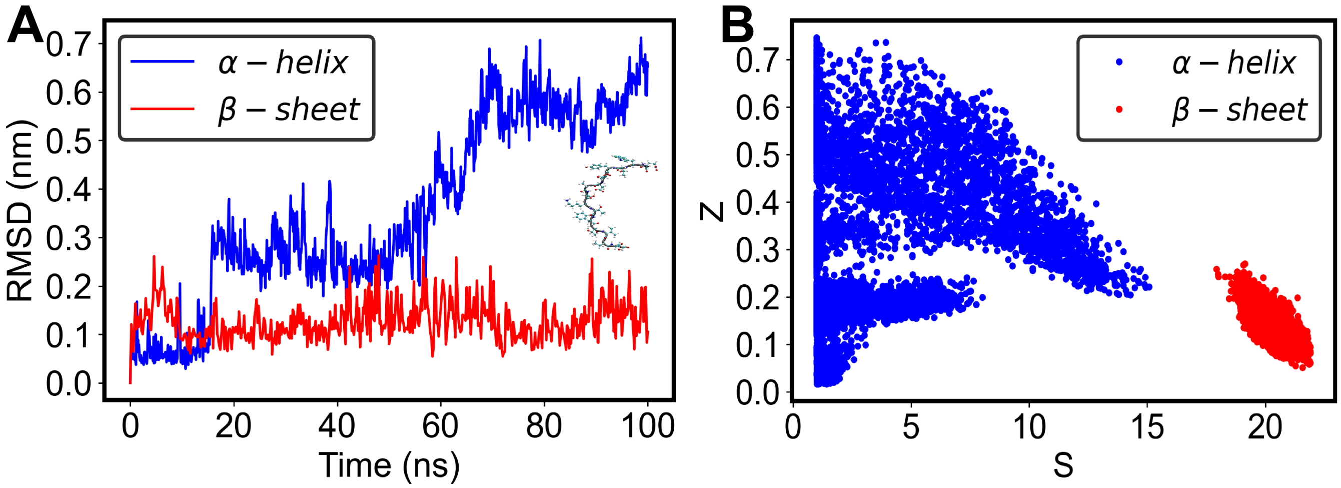


Figure 3: **G-protein unbiased simulations** (A) RMSD from the unbiased simulations of $\alpha$-helix and $\beta$-hairpin structures. $\alpha$-helix is completely unfolded after 60 ns of simulation. Inset illustrates the final structure of the $\alpha$-helix at the end of the simulation. (B) $S$ and $Z$ sampling in unbiased simulations of $\alpha$-helix and $\beta$-hairpin states.

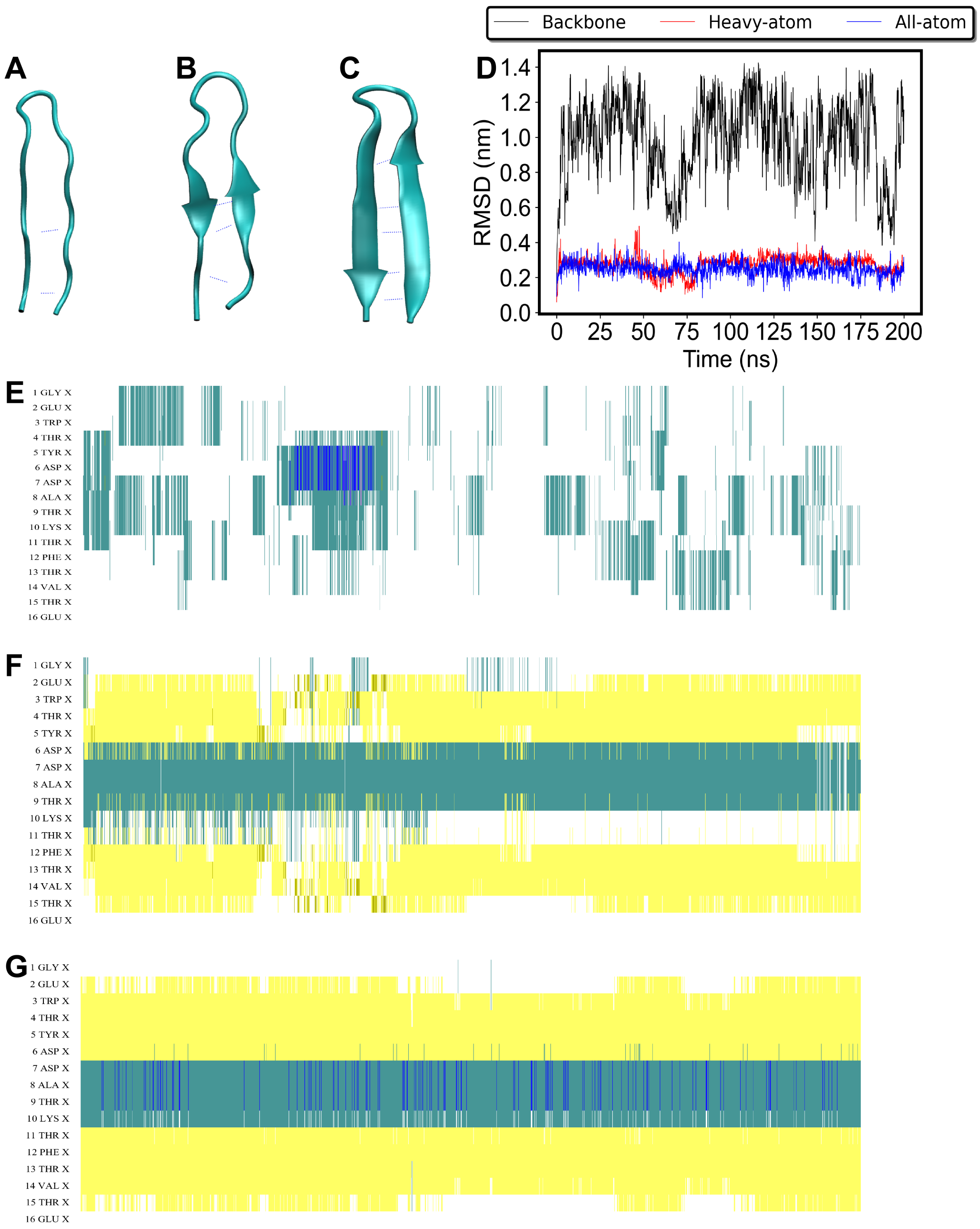


Figure 4: **G-protein pulling simulations** Unbiased simulation of the $\beta$-hairpin obtained from the pulling simulations from $\alpha$-helix to $\beta$-hairpin in $S$ and $Z$ space with 22 reference states for the reference state having (A) backbone atom information in the reference path, final pulled state ($S, Z$ = 21.99, 0.011) has 2 Hbonds, (B) heavy atom information in the reference path, final pulled state ($S, Z$ = 21.93, 0.021) has 3 Hbonds, and (C) all atom information in the reference path, final pulled state ($S, Z$ = 21.99, 0.013) has 5 Hbonds. (D) RMSD of the unbiased simulations of the final pulled $\beta$-hairpin state indicates that only the backbone atom information in the reference path is insufficient to form a stable $\beta$-hairpin. Changes in the secondary structure with time from unbiased simulation for (E) backbone atom information, (F) heavy atom information, and (F) all atom information in the reference path, indicate that including the all atom information in the reference path stabilizes the $\beta$-hairpin to the greatest extent.

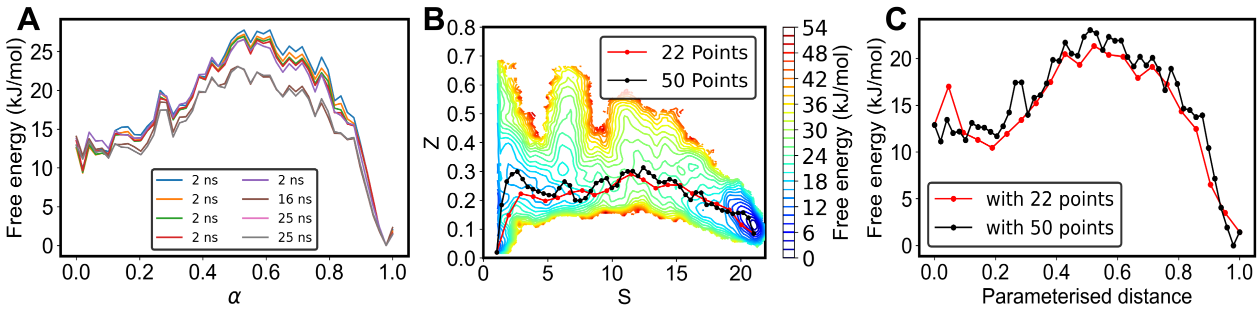


Figure 5: **G-protein free energy estimation** (A) Convergence of the free energy profile projected along the converged path, calculated at every additional sampling with last three simulations performed for 16 ns, 25 ns, and 25 ns on each points on the converged path. (D) Location of the converged string defined by 22 and 50 points. (E) Free energy profile along the 22 point string and 50 point string. Free energy barrier is 21.35 kJ/mol for 22 points string, and 23.03 kJ/mol for 50 points string. The difference is within $\sim k_B T$.
